# Quantitative Morphology of Stump Sprouts Reveals a Common Growth Axis and a Reproducible Basal Longitudinal Cavity in an Ornamental Cherry

**DOI:** 10.64898/2026.09.06.749736

**Authors:** Nagi Nangirky Ogata, Norichika Ogata

## Abstract

Stump sprouts may originate from pre-existing preventitious buds or adventitious buds, but whether these developmental origins produce distinguishable whole-shoot phenotypes remains unclear. We examined all visible sprouts on a single stump of an ornamental cherry (Prunus sp.) and measured four preselected traits: fresh weight, basal diameter, shoot length, and node number. The initial census comprised 35 sprouts. These traits were strongly correlated and were dominated by a common multivariate growth axis. Unsupervised mixture modelling identified statistical components within the morphological distribution, but these components could not be assigned to preventitious or adventitious developmental origins. During subsequent examination, we identified an unexpected basal longitudinal cavity and recorded its presence and axial length. The cavity occurred in 23 of 31 sectioned sprouts (74%). After complete removal of the initial sprout population, all visible sprouts were collected again seven days later. This second cohort comprised 22 substantially smaller sprouts but reproduced the coordinated whole-shoot growth structure. Remarkably, the basal longitudinal cavity occurred in 16 of 22 sprouts (73%), nearly identical to its initial frequency despite the marked difference in shoot size. Cavity presence was largely independent of overall shoot size, whereas cavity length increased with size among cavity-positive sprouts. Thus, gross quantitative morphology did not discriminate the hypothesized developmental origins, whereas basal anatomy revealed a reproducible feature potentially informative of developmental history. Direct anatomical and ontogenetic analyses are required to determine whether the basal longitudinal cavity is associated with sprout origin.

## Introduction

Stump sprouting is an important mode of vegetative regeneration in many woody plants. Shoots emerging from stems, stumps, or root collars may arise from buds of different developmental origins, but terminology for these structures has historically been inconsistent. Stone and Stone (1943) emphasized that *adventitious* should refer to developmental origin rather than simply to the appearance of a shoot following injury. They distinguished dormant or latent buds formed within the normal phyllotactic system from adventitious buds arising outside that system, while explicitly noting that an adventitious bud, once formed, may itself remain dormant. Thus, dormancy and developmental origin are distinct properties of a bud (Stone and Stone, 1943).

Anatomical studies subsequently provided a structural basis for this distinction. Fink (1983) described endogenous adventitious buds arising from parenchymatous tissues in the bark of several temperate and tropical trees, while also showing that originally exogenous dormant buds may become embedded by secondary growth and consequently resemble endogenous buds. This observation is important because the apparent position of a bud or shoot within mature tissues does not necessarily reveal its developmental origin. In *Quercus petraea*, Fontaine et al. (1998) similarly defined proventitious epicormic buds as derivatives of pre-existing buds within the normal leaf- or scale-axil system and connected by vascular tissues toward the main-stem pith. They further showed that these bud systems are developmentally dynamic: secondary bud primordia may develop in association with persistent primary epicormic buds, maintaining or increasing a reservoir of potential shoots over time.

Such persistent bud systems have also been demonstrated directly in stump-sprouting species. In *Betula pubescens*, dormant basal buds originate as axillary buds during the seedling stage, maintain their vascular connections during subsequent radial growth, and may branch to form clusters of secondary buds (Kauppi et al., 1987). Felling causes many of these pre-existing basal buds to burst, although only a subset ultimately develops into sprouts. In cutting-origin stools of *Salix* ‘Aquatica’, primary axillary buds are accompanied by collateral accessory buds; the primary bud generally bursts first, whereas accessory buds may remain inactive and burst later, sometimes after several years. Coppicing strongly accelerates their development into shoots (Paukkonen et al., 1992). Consequently, sequential cohorts of shoots following disturbance do not necessarily require sequential de novo formation of adventitious buds.

Classical studies of stump sprouting provide complementary evidence from wood anatomy. Roth and Hepting (1943) showed that oak stump sprouts commonly arise from latent buds that maintain radial wood connections toward the primary xylem as the stem increases in diameter. Such connections leave bud traces through the wood and provide anatomical evidence of pre-existing bud origin. Woods and Cassady (1961) subsequently demonstrated experimentally that developmental origin can change during repeated sprouting. Cross-sections of scrub-oak stumps showed that the initial sprouts following cutting originated from dormant buds, as indicated by radial traces extending from the pith to the cambium. After these sprouts were removed, callus formed over injured tissues and became the source of numerous new buds and sprouts. The authors concluded that the initial stump sprouts were entirely derived from dormant buds, whereas succeeding sprouts after removal were nearly all derived from adventitious buds. Thus, sprouts of different developmental origins can occur during the regenerative history of a single stump.

The distinction has also been incorporated into silvicultural classifications. De Simón and Bocio (1999) distinguished proventitious sprouts arising from dormant buds already present in living tissues of the stump base, root collar, or major roots from adventitious sprouts associated with wound healing and callus formation. More recently, Ríos-Saucedo et al. (2017) quantified sprouting in several woody species while distinguishing proventitious and adventitious shoots. In that study, however, shoot type was assigned from physiological characteristics, position, and attachment to the stump rather than inferred from quantitative shoot phenotype.

Quantitative morphology has nevertheless been used extensively to characterize stump sprouts and related epicormic formations. Measurements such as shoot number, length, diameter, biomass, and branching have been used to quantify sprouting vigor and regeneration. Fontaine et al. (2004) went further by quantitatively classifying epicormic formations of *Q. petraea* according to external morphology and relating those formations to traces and defects within the wood. Their classification distinguished thin shoots, shoot stumps, bud clusters, and thicker persistent epicormic shoots, and proposed ontogenetic transitions among these structures. Their review of previous work also noted earlier quantitative classifications based on the number and length of visible epicormic shoots. Thus, quantitative morphology can discriminate developmental or structural states within an epicormic system.

These studies raise a different question: is developmental origin itself expressed strongly enough in the resulting shoot phenotype to be detected without first assigning an origin to individual shoots? If preventitious and adventitious shoots differ systematically in establishment, vascular connection, or access to resources in the parent stump, their distinct developmental histories might generate separable multivariate phenotypic populations. Conversely, once a shoot apical meristem becomes active, subsequent shoot development might converge onto a common growth program and obscure information about bud origin. The latter possibility is consistent with the developmental anatomy described by Fontaine et al. (1998): proventitious epicormic buds of *Q. petraea* contain no preformed shoot, and the emerging shoot is instead neoformed by the terminal meristem during emergence. Similar organization has been described for inhibited buds responsible for stump sprouting in other woody plants. Although quantitative morphology has previously been used to characterize stump sprouts and classify epicormic formations, we found no previous study that directly tested whether preventitious and adventitious developmental origins can be recovered as intrinsic population structure from the multivariate whole-shoot phenotype of unlabeled stump sprouts.

We addressed this question using a single stump of an ornamental cherry (*Prunus* sp.) growing near Tama-Plaza Station, Yokohama, Japan. Rather than subsampling the sprout population, we removed every visible sprout present on the stump on 24 August 2026. Before collection, four quantitative traits—fresh weight, basal diameter, shoot length, and number of nodes—were selected to characterize the whole-shoot phenotype. By examining all sprouts arising from a single stump, variation among source trees was inherently excluded. We asked whether the resulting multivariate phenotype contained discrete population structure potentially consistent with heterogeneous developmental origins.

Inspection of the collected material subsequently revealed variation at the shoot base that had not been included in the original measurements. Some sprouts retained attached woody tissue whereas others did not. Longitudinal sectioning further revealed a macroscopically visible cavity extending from the basal end of many sprouts. We refer to this feature descriptively as the **basal longitudinal cavity** (BLC), without assuming its tissue identity, developmental origin, or function.

Developmentally organized internal spaces are known from woody buds and shoots. Sakai (1979), for example, demonstrated a freezing-avoidance system in conifer primordial shoots in which water migrates out of the primordial shoot and freezes outside the sensitive tissues; masses of ice accumulated mainly beneath the crown. Related variation in freezing anatomy occurs within *Prunus*: Kadir and Proebsting (1994) examined flower buds of 20 *Prunus* species and found markedly different freezing and supercooling strategies among species. These structures and phenomena are not evidence that the cavity observed in the present stump sprouts is homologous to a bud cavity or performs a freezing-related function; rather, they establish that developmentally organized internal spaces and water compartments occur in woody buds and shoots.

A second possibility is secondary tissue loss associated with decay. Roth and Hepting (1943) demonstrated that decay in a parent oak stump can progress into an attached sprout through their woody connection. Long-term observations of baldcypress stump sprouts likewise found that rot from old stumps often appeared to spread into sprout bases (Keim et al., 2006). The latter observations concerned sprouts on stumps harvested 10–41 years previously and therefore represent a very different developmental timescale from newly emerging sprouts. We consequently treated the presence and longitudinal extent of the basal cavity as descriptive anatomical variables rather than evidence for either a developmental or decay-related mechanism.

Complete removal of the first visible sprout population also constituted a perturbation of the stump-sprouting system. We therefore returned to the same stump seven days later, on 31 August 2026, and again collected every visible sprout. This second census was particularly informative in light of the classical observation that removal of initial dormant-bud-derived sprouts can be followed by adventitious sprouting from callus (Woods and Cassady, 1961). However, emergence following removal does not itself establish developmental origin. Stone and Stone (1943) explicitly noted that an adventitious bud, once formed, may remain dormant, whereas the studies of *Betula, Salix*, and *Quercus* demonstrate that pre-existing bud systems may contain secondary or accessory buds capable of later development (Kauppi et al., 1987; Paukkonen et al., 1992; Fontaine et al., 1998). The second cohort could therefore have included shoots from previously inactive preventitious buds, previously formed but dormant adventitious buds, buds or shoots already developing but externally invisible at the first census, or adventitious buds newly formed following removal.

We therefore used two complete censuses of the same stump to address two related questions: (i) whether multivariate whole-shoot phenotype contains statistical population structure that can be related to the hypothesized preventitious and adventitious developmental origins, and (ii) whether the phenotypic structure and newly recognized basal anatomical characteristics recur in a new cohort following complete removal of the initially visible sprouts.

## Materials and Methods

### Study site and plant material

The study was conducted on a stump of an ornamental cherry (*Prunus* sp.) growing near Tama-Plaza Station, Yokohama, Japan. The species and cultivar of the tree were not determined. The stump was old and partially hollow, with visible deterioration of the central portion. Numerous shoots were present around the stump margin and basal region. Individual shoots arising from the stump are referred to here as stump sprouts.

The study was designed as a complete census of the visible sprout population of a single stump rather than as a subsampling experiment. Consequently, all visible sprouts present at each census were collected. This design excluded variation among source trees from comparisons among sprouts, although the single-stump design does not provide replication among trees.

### First complete census and quantitative morphological measurements

The first complete census was conducted on 24 August 2026. Before collection, four quantitative traits were selected to characterize whole-shoot morphology: fresh weight, basal diameter, shoot length, and number of nodes. The initial question was whether variation in these quantitative traits could reveal distinct morphological populations potentially corresponding to preventitious and adventitious developmental origins.

All visible sprouts on the stump were removed, yielding 35 specimens. Fresh weight was measured in 0.001 grams using an analytical balance LA-JT1003D(LA) (Lachoi Scientific Instruments Co. Ltd., Zhejiang, China). Basal diameter was measured in 0.1 millimeters near the base of each sprout using a caliper SLIDE RULER (Designphil Inc., Tokyo, JAPAN), and shoot length was measured in centimeters from the basal end to the shoot apex using a 50-cm gridded ruler (STAEDTLER 062 06-50, STAEDTLER, Germany). Node number was determined by direct counting. These four measurements constituted the preselected quantitative morphological dataset.

The developmental origin of individual sprouts was not assigned during collection. In particular, sprouts were not classified as preventitious or adventitious according to their external position or gross morphology because the purpose of the analysis was to determine whether developmental heterogeneity could be recovered from the quantitative phenotype itself.

### Examination of basal features

During examination of the collected specimens after the first census, some sprouts were observed to retain pieces of woody tissue at their basal ends whereas others did not. The presence or absence of attached woody tissue was therefore recorded as an additional qualitative character. Because this character was recognized after collection, it was treated as an exploratory rather than a preselected variable.

Subsequent longitudinal sectioning of the sprout bases revealed a macroscopically visible internal cavity extending longitudinally from the basal end in some specimens. This feature was designated the basal longitudinal cavity (BLC). The term is used descriptively and does not imply that the cavity represents pith, decay, or any particular developmental structure. BLC presence or absence was recorded, and, when present, its axial length was measured in millimeters from the basal end along the longitudinal axis of the cavity.

Thirty-one of the 35 sprouts collected on 24 August were longitudinally sectioned and examined for the BLC. Four specimens were retained intact and were therefore treated as missing, rather than BLC-negative, for analyses of cavity occurrence. Thus, BLC prevalence in the first census was calculated from the 31 sectioned specimens.

### Complete removal and second census

Collection of all visible sprouts on 24 August resulted in complete removal of the externally visible sprout population and thereby constituted a perturbation of the stump. The stump was examined again seven days later, on 31 August 2026. All sprouts visible at that time were again collected, yielding a second complete census of 22 specimens.

The same four quantitative morphological traits—fresh weight, basal diameter, shoot length, and node number—were measured for all 22 sprouts using the same procedures as in the first census. Attached woody tissue was recorded for each specimen. All 22 sprouts were longitudinally sectioned and examined for BLC presence and, when present, BLC axial length was measured using a caliper SLIDE RULER (Designphil Inc., Tokyo, JAPAN). In total, the two censuses yielded 57 sprouts for quantitative morphological analysis and 53 longitudinally sectioned sprouts for BLC analysis.

The seven-day interval was defined according to the time between the two complete censuses and was not assumed to represent the age of the underlying buds. Sprouts observed in the second census could therefore have originated from developmental structures established either before or after the first census. Raw data for all 57 sprouts are provided in Table 1.

**Table 1.**
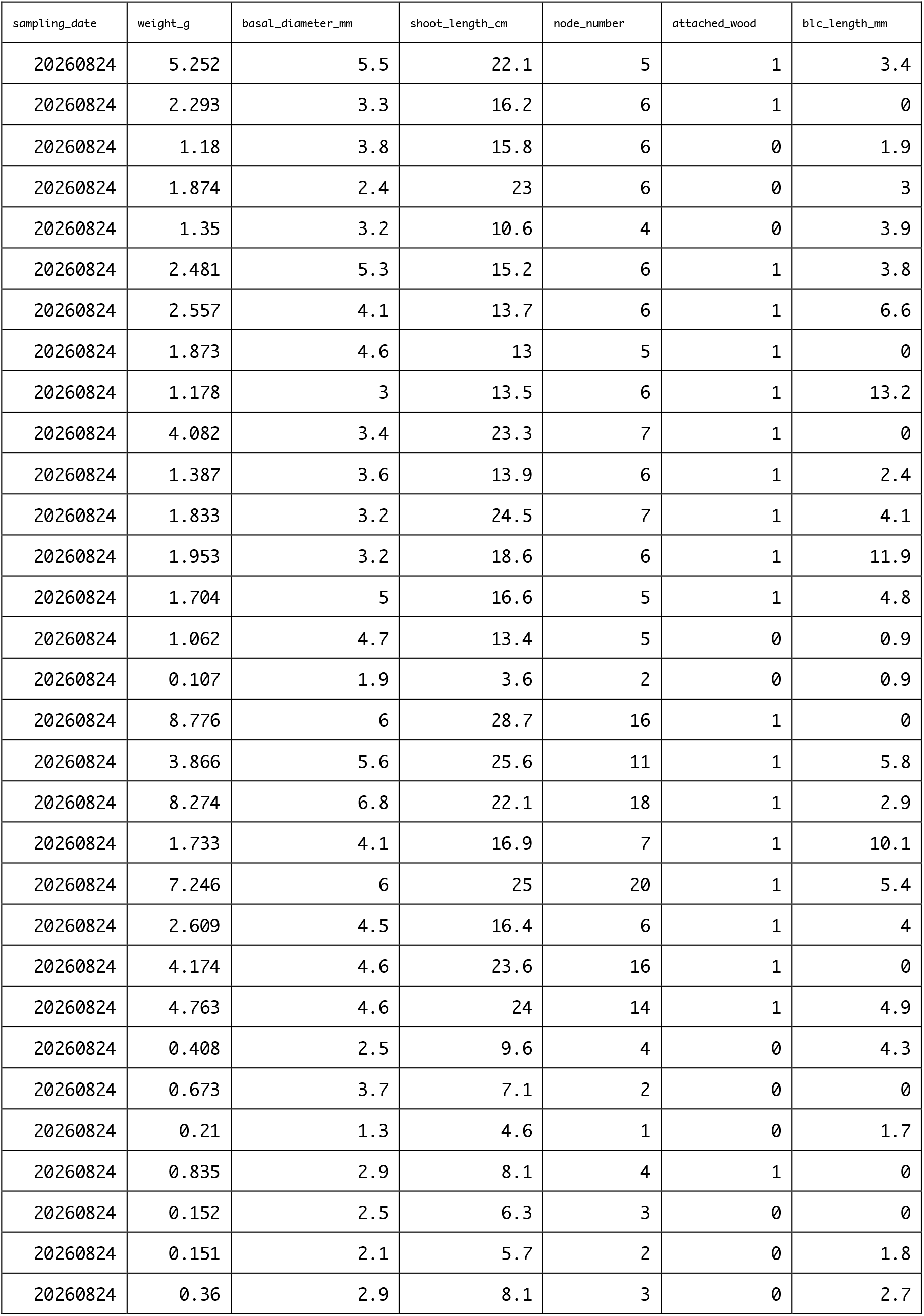
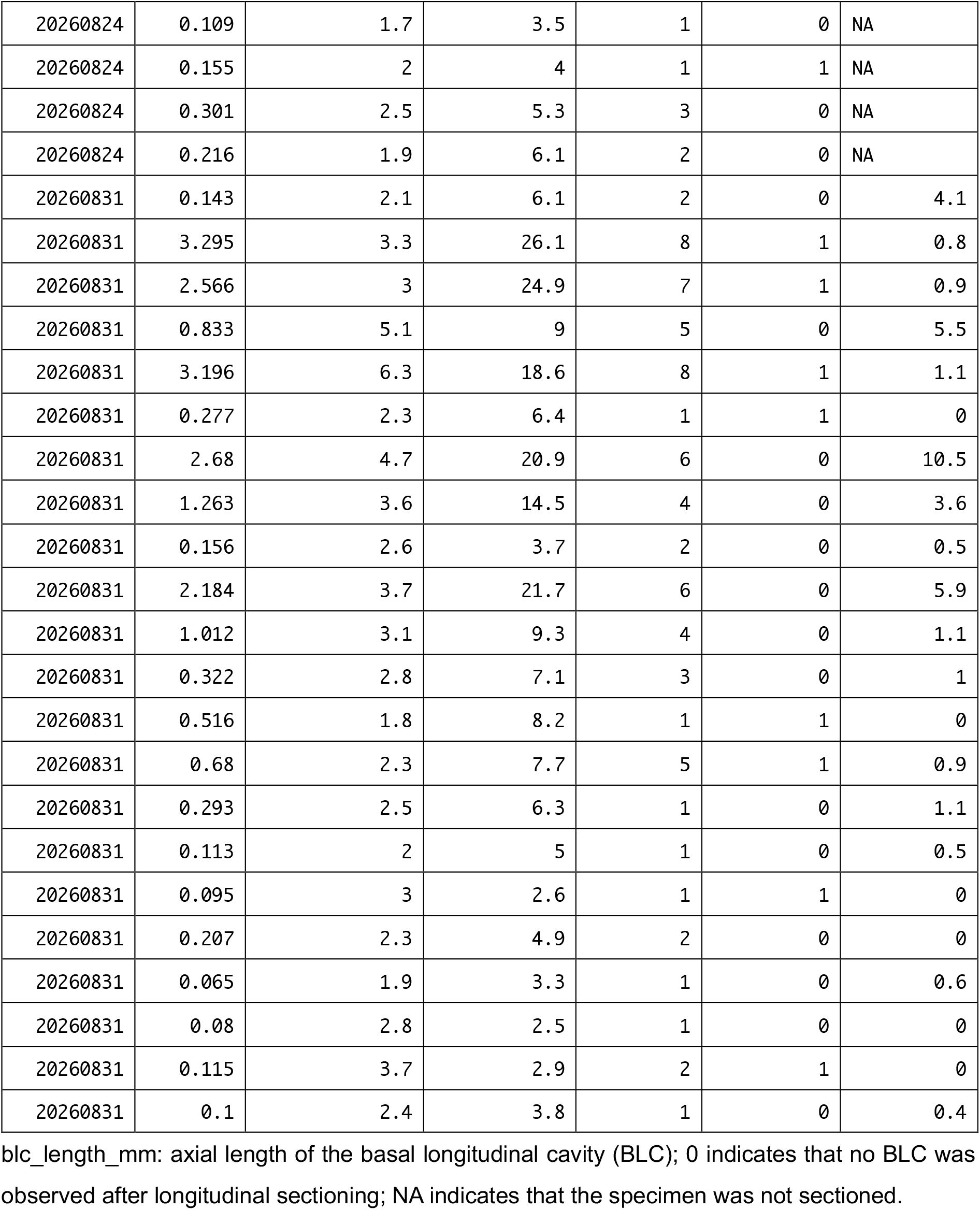
Raw morphological and basal anatomical measurements of stump sprouts collected in the two complete censuses.

### Multivariate analysis of quantitative morphology

Fresh weight, basal diameter, shoot length, and node number were analyzed as quantitative whole-shoot traits. Pairwise associations among the four variables were evaluated using Spearman’s rank correlation coefficient (ρ).

For multivariate analysis, each of the four traits was standardized to zero mean and unit variance across the pooled dataset of 57 sprouts. Principal component analysis (PCA) was then performed on the standardized variables. The first principal component (PC1) was interpreted according to its loadings and used as a summary measure of coordinated whole-shoot variation. Because all four traits contributed in the same direction, PC1 represented an overall growth-size axis.

The two census cohorts were visualized in the common PCA space. PC1 scores were compared between the 24 August and 31 August cohorts using the Wilcoxon rank-sum test. Because GMM represents the observed distribution as a mixture of Gaussian component distributions, departure from Gaussianity was quantified for each quantitative trait using quantile–quantile root mean square error (QQ-RMSE), calculated as the root mean square deviation between standardized observed quantiles and the corresponding theoretical standard-normal quantiles (Ogata, 2026). QQ-RMSE was calculated before and after natural-log transformation. Because log transformation reduced QQ-RMSE for all four traits, the log-transformed values were standardized and used for subsequent GMM analysis. Gaussian mixture models (GMMs) were fitted using Python 3.13.5 and scikit-learn 1.8.0 (`sklearn.mixture.GaussianMixture`). Models containing one to three Gaussian components were fitted using unrestricted component-specific covariance matrices (`covariance_type=“full”`) and compared using the Bayesian information criterion (BIC), with the model having the lowest BIC considered the best-supported among those examined. GMM components were treated as statistical partitions of the morphological distribution and were not assumed *a priori* to correspond to preventitious or adventitious developmental origins.

### Analysis of attached woody tissue and the basal longitudinal cavity

The frequency of attached woody tissue was compared between the two censuses using Fisher’s exact test. Within the initial census, quantitative morphological traits were compared between sprouts with and without attached woody tissue using Mann–Whitney tests. The association between attached woody tissue and the common multivariate growth axis was additionally examined by logistic regression using PC1 as the explanatory variable.

BLC prevalence was calculated separately for each census using only longitudinally sectioned specimens and was compared between censuses using Fisher’s exact test. BLC axial length was analyzed separately from BLC presence. Among BLC-positive sprouts, BLC lengths were compared between census dates using the Wilcoxon rank-sum test.

To examine whether BLC extension was associated with overall sprout growth, the relationship between BLC axial length and pooled PC1 was evaluated among BLC-positive sprouts using Spearman’s rank correlation. Correlations were calculated separately for the 24 August and 31 August cohorts and for the pooled BLC-positive dataset. Linear regression lines shown in graphical presentations were used only as descriptive visual summaries; statistical inference for the association between PC1 and BLC length was based on Spearman’s rank correlation.

### Statistical analysis

Most statistical analyses were performed in R version 4.4.1. Tests were two-sided, and *P* < 0.05 was considered statistically significant. Unless otherwise specified, nonparametric tests were used for comparisons of continuous variables because of the small sample sizes and the strongly unequal distributions of sprout size. Missing observations were excluded only from analyses requiring the corresponding variable; specifically, the four intact specimens from the first census were excluded from BLC analyses but retained in analyses of whole-shoot morphology and attached woody tissue.

## Results

### Complete census and re-emergence of stump sprouts after removal

At the first census on 24 August 2026, all 35 visible sprouts arising from the stump were collected (Fig. 1A, B). Fresh weight, basal diameter, shoot length, and node number were measured for all sprouts. These four quantitative traits had been selected before the first census. During subsequent examination of the collected specimens, the presence or absence of attached woody tissue was additionally recorded. Longitudinal sectioning further revealed a previously unrecorded basal feature, here termed the basal longitudinal cavity (BLC) (Fig. 1C). Thirty-one of the 35 sprouts were longitudinally sectioned and examined for BLC presence and axial length; four specimens were retained intact and were not sectioned. The representative specimen in Fig. 1C shows the BLC extending longitudinally from the basal end of the sprout.

**Figure 1.**
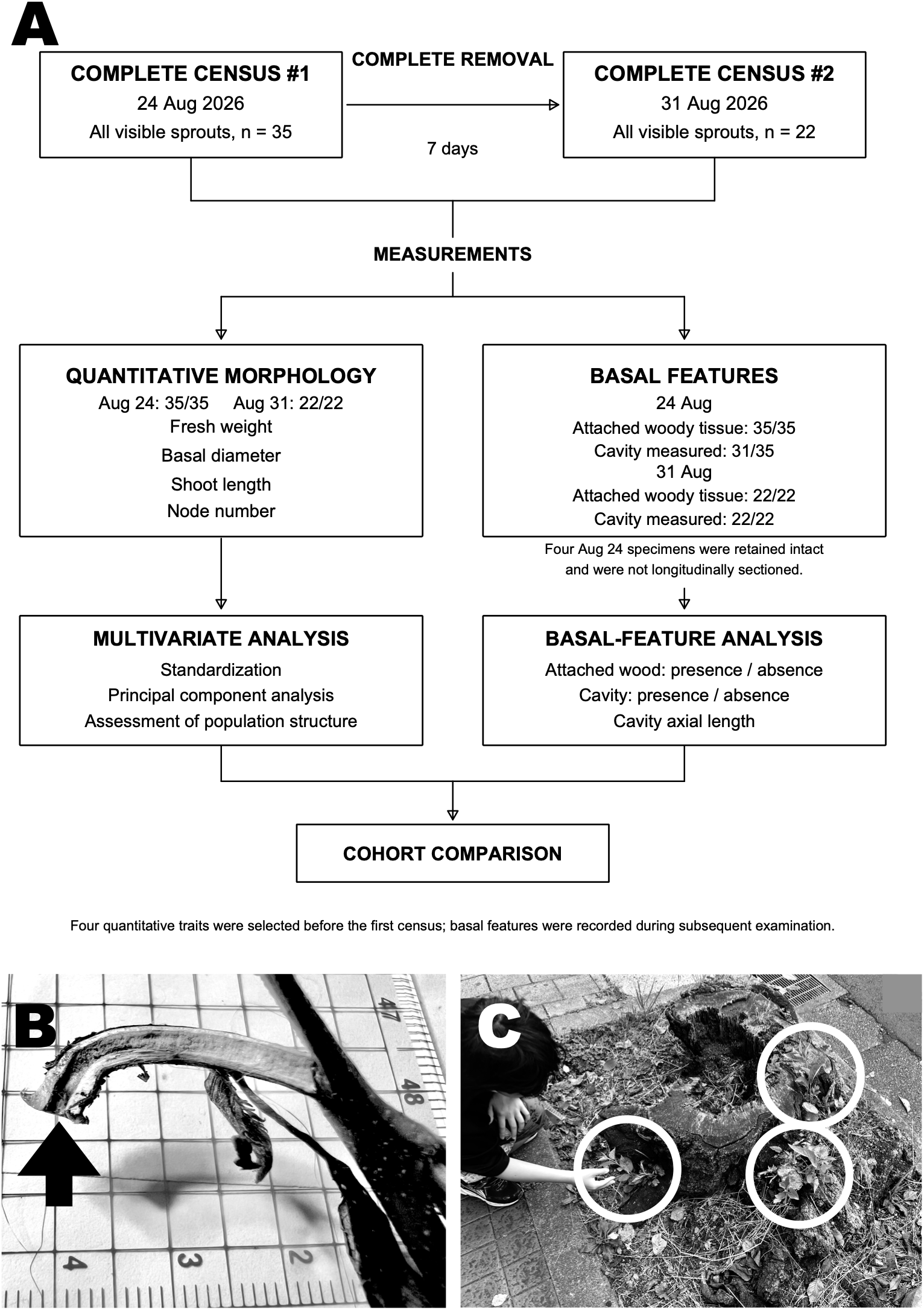
Study design, study stump, and basal longitudinal cavity in stump sprouts of an ornamental cherry (Prunus sp.). **(A)** Study design. All visible sprouts on the stump were collected on 24 August 2026 (n = 35), followed by complete removal of the visible sprout population. Seven days later, on 31 August 2026, all newly visible sprouts were collected again (n = 22). Fresh weight, basal diameter, shoot length, and node number were measured for all 57 sprouts. Attached woody tissue was also recorded for all sprouts. Thirty-one of the 35 sprouts from the first census and all 22 sprouts from the second census were longitudinally sectioned and scored for the presence and axial length of a basal longitudinal cavity (BLC); four specimens from the first census were retained intact. The four quantitative morphological traits were selected before the first census, whereas attached woody tissue and BLC were recorded during subsequent examination. **(B)** Study stump of an ornamental cherry (*Prunus* sp.). White circles indicate regions bearing stump sprouts. **(C)** Representative longitudinal section of a sprout base showing the BLC (arrow). The specimen was photographed on a grid with 5-mm spacing, which provides a dimensional reference for the image.

The first census was followed by complete removal of all visible sprouts from the stump. Seven days later, on 31 August 2026, 22 visible sprouts were present. All 22 were collected, measured for the same four quantitative morphological traits, scored for attached woody tissue, and longitudinally sectioned for examination of the BLC (Fig. 1A). Thus, the two complete censuses yielded 57 sprouts in total: 35 in the initial census and 22 in the second census seven days after complete removal. The study design and numbers of specimens subjected to each analysis are summarized in Fig. 1A.

### Quantitative whole-shoot morphology was dominated by a common growth axis

The four preselected quantitative traits were strongly correlated with one another (Fig. 2A). Principal component analysis of the standardized measurements showed that most multivariate variation was concentrated along a single axis (Fig. 2B). PC1 accounted for **85.0%** of the total variance, whereas PC2 accounted for **8.1%**. All four traits contributed positively to PC1, identifying it as an overall growth-size axis. Sprouts collected on 24 and 31 August overlapped extensively in PC1–PC2 space, although the second cohort occupied predominantly the smaller end of this common growth axis. PC1 scores were significantly lower on 31 August than on 24 August (Fig. 2C; Wilcoxon rank-sum test, *P* = 0.031).

**Figure 2.**
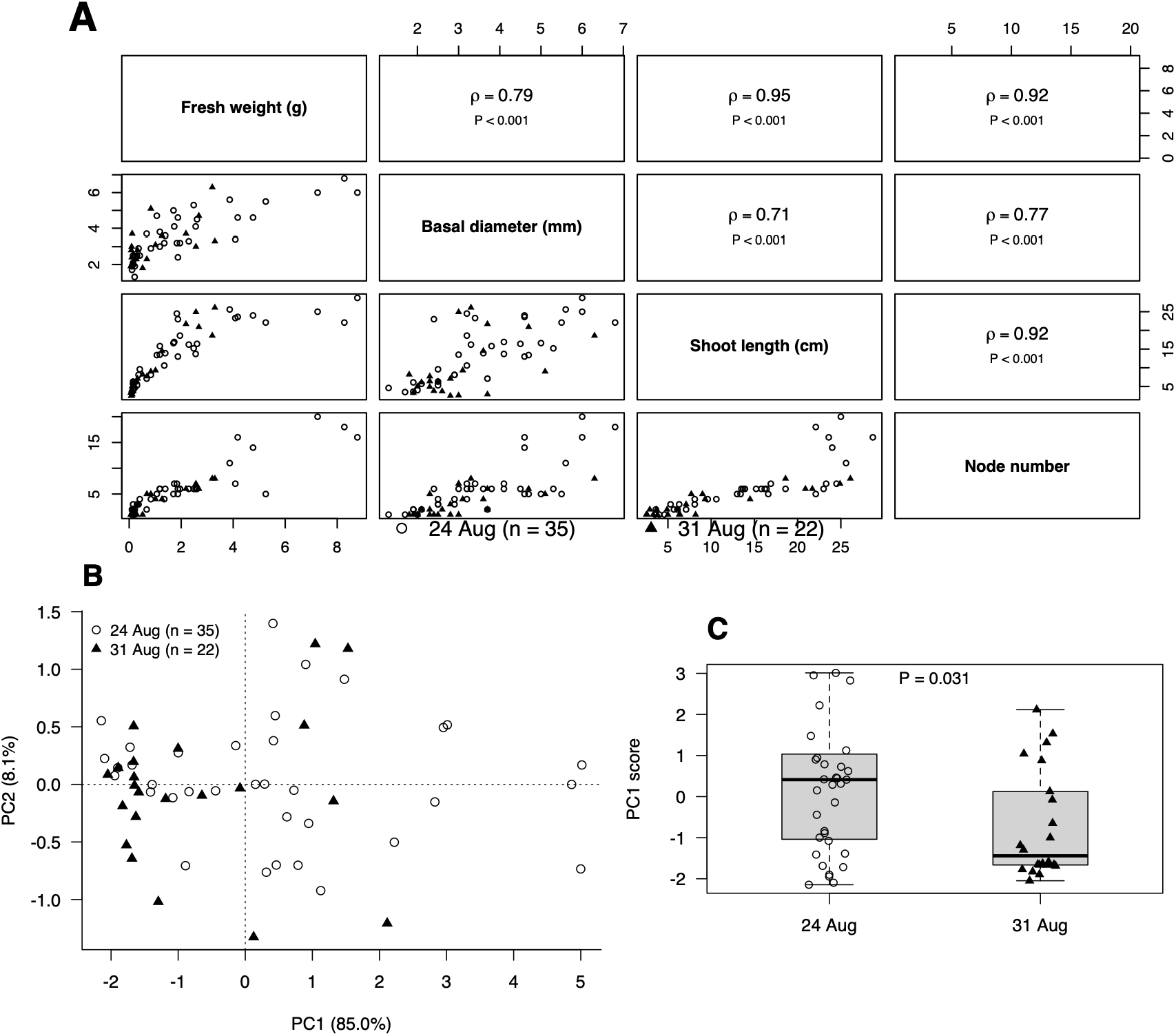
Quantitative whole-shoot morphology was dominated by a common growth axis across two successive stump-sprout cohorts. **(A)** Pairwise relationships among the four preselected quantitative traits: fresh weight, basal diameter, shoot length, and node number. Individual sprouts collected on 24 August (n = 35) and 31 August (n = 22) are shown by open circles and filled triangles, respectively. Spearman’s rank correlation coefficients (ρ) and associated *P* values are shown for each pair of traits. **(B)** Principal component analysis (PCA) of the four standardized quantitative traits for all 57 sprouts. PC1 accounted for 85.0% of the total variance and represented a common growth-size axis. Sprouts from the two censuses overlapped extensively along the common multivariate growth axis rather than forming distinct cohort-specific groups. **(C)** Distribution of PC1 scores in the two censuses. Individual observations are shown using the same symbols as in panel B, with boxplots indicating the median and interquartile range. PC1 scores were compared between censuses using a Wilcoxon rank-sum test.

The raw quantitative traits showed varying degrees of departure from Gaussianity, with QQ-RMSE values of 0.4880, 0.2324, 0.2866, and 0.4558 for fresh weight, basal diameter, shoot length, and node number, respectively. Natural-log transformation reduced the corresponding values to 0.2234, 0.1173, 0.2386, and 0.2629, respectively, and the transformed values were therefore used for GMM analysis. Gaussian mixture modelling of the log-transformed standardized traits favored a three-component solution. BIC values were 387.46, 415.96, and 327.18 for the one-, two-, and three-component models, respectively. The three components contained 11, 29, and 17 sprouts, respectively, and were ordered predominantly along the common growth-size axis. The smallest component consisted entirely of one-node sprouts, whereas the largest component comprised larger, more developed shoots. All three components contained sprouts from both sampling cohorts.

Thus, although the sprouts present seven days after complete removal were smaller overall, their quantitative morphology remained embedded within the same coordinated multivariate growth continuum observed in the initial census. Unsupervised mixture modelling identified statistical components within this continuum, but these components primarily reflected differences along the growth-size axis rather than census-specific groups.

### Basal longitudinal cavity prevalence was reproduced in the second cohort

The BLC was present in **23 of 31 sectioned sprouts (74.2%)** in the initial census and absent in eight (Fig. 3A). In the second census, the BLC was present in **16 of 22 sprouts (72.7%)** and absent in six. Thus, despite complete removal of the visible sprout population seven days earlier and the smaller overall size of the second cohort, BLC prevalence was nearly identical between the two censuses (Fisher’s exact test, *P* = 1.0).

**Figure 3.**
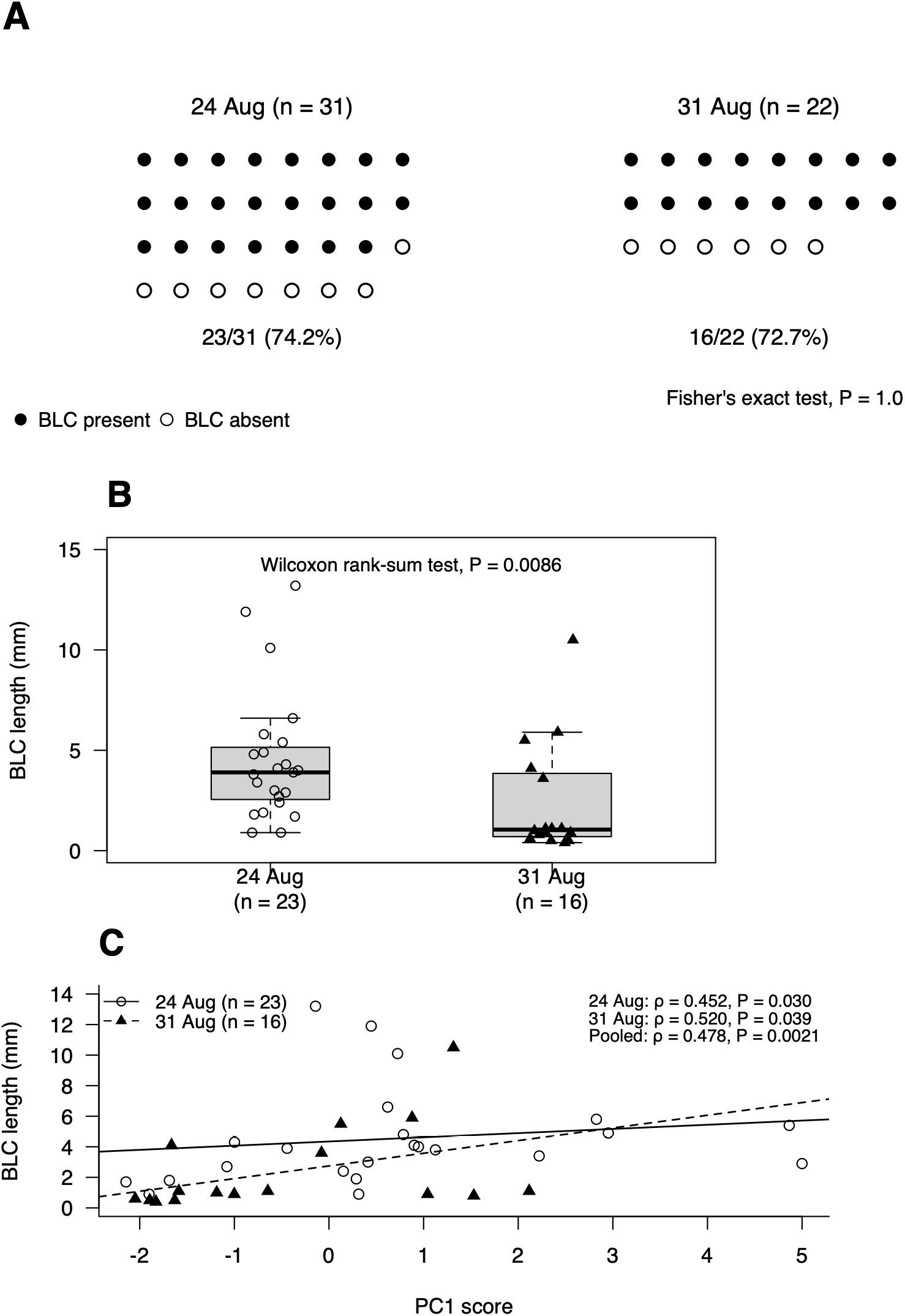
Occurrence and axial length of the basal longitudinal cavity in two successive stump-sprout cohorts. **(A)** Occurrence of the basal longitudinal cavity (BLC) in longitudinally sectioned sprouts from the two complete censuses. Each symbol represents one sprout; filled and open circles indicate BLC presence and absence, respectively. The BLC was present in 23 of 31 sprouts (74.2%) on 24 August and 16 of 22 sprouts (72.7%) on 31 August. BLC prevalence did not differ between censuses (Fisher’s exact test, *P* = 1.0). **(B)** BLC axial length among BLC-positive sprouts. Individual sprouts are shown as open circles for 24 August (n = 23) and filled triangles for 31 August (n = 16); boxplots show the median and interquartile range. BLCs were shorter in the second cohort (median, 1.05 mm) than in the initial cohort (median, 3.9 mm; Wilcoxon rank-sum test, *P* = 0.0086). **(C)** Relationship between BLC axial length and the pooled PC1 score among BLC-positive sprouts. PC1 was calculated from the four standardized quantitative morphological traits for all 57 sprouts. BLC length was positively correlated with PC1 in both the 24 August cohort (Spearman’s ρ = 0.452, *P* = 0.030, n = 23) and the 31 August cohort (ρ = 0.520, *P* = 0.039, n = 16), as well as in the pooled data (ρ = 0.478, *P* = 0.0021, n = 39). Solid and dashed lines indicate descriptive linear fits for the 24 August and 31 August cohorts, respectively.

In contrast to its prevalence, the axial length of the BLC differed between cohorts (Fig. 3B). Among BLC-positive sprouts, median BLC length was **3.9 mm** in the initial cohort (n = 23) and **1.05 mm** in the second cohort (n = 16). BLCs were significantly shorter in the second cohort (Wilcoxon rank-sum test, *P* = 0.0086).

BLC length was also associated with the multivariate growth-size axis (Fig. 3C). Among BLC-positive sprouts, BLC length was positively correlated with pooled PC1 within the initial cohort (Spearman’s ρ = 0.452, *P* = 0.030, n = 23) and within the second cohort (ρ = 0.520, *P* = 0.039, n = 16). The relationship remained significant when both cohorts were pooled (ρ = 0.478, *P* = 0.0021, n = 39). Thus, larger sprouts tended to have longer BLCs within each cohort, whereas the probability of BLC occurrence itself remained nearly unchanged between two cohorts that differed substantially in overall shoot size.

### Attached woody tissue was associated with sprout size

Attached woody tissue was observed in 21 of 35 sprouts (60.0%) in the initial census and 8 of 22 sprouts (36.4%) in the second census. The difference in frequency between censuses was not statistically significant (Fisher’s exact test, *P* = 0.106).

Within the initial census, sprouts with attached woody tissue were larger than those without it for all four quantitative traits (Mann–Whitney tests: fresh weight, *P* = 2.76 × 10^−5^; basal diameter, *P* = 3.53 × 10^−4^; shoot length, *P* = 1.85 × 10^−4^; node number, *P* = 9.02 × 10^−5^). Consistent with these univariate relationships, the probability of attached woody tissue increased with PC1 (logistic regression coefficient = 1.15, *P* = 0.002; odds ratio ≈ 3.17 per unit increase in PC1). Attached woody tissue therefore varied strongly with the overall growth-size axis rather than defining an obviously independent class of sprouts.

## Discussion

The present study asked whether developmental heterogeneity among stump sprouts could be recovered from quantitative whole-shoot morphology. The four preselected traits were strongly correlated, and PC1 alone accounted for 85.0% of the total multivariate variation, representing a common axis of overall shoot size or developmental stage. Although GMM identified three statistical components, their ordering along the common growth-size axis, the restriction of the smallest component to one-node sprouts, and the occurrence of all three components in both cohorts suggest that the mixture structure is more parsimoniously interpreted as partitioning of developmental size variation than as evidence of independently identifiable developmental-origin classes. Thus, quantitative whole-shoot morphology contained statistical structure, but this structure did not provide a basis for distinguishing the hypothesized preventitious and adventitious developmental origins. A second complete census, performed seven days after removal of all initially visible sprouts, yielded substantially smaller shoots but reproduced the same strongly coordinated morphological structure. Thus, within the resolution of the traits examined here, developmental heterogeneity, if present, was not expressed as distinct whole-shoot phenotypic populations.

This result should not be interpreted as evidence that all sprouts had the same developmental origin. Classical anatomical studies indicate that developmental origin can be remarkably difficult to infer from the final position or structure of a bud. Fink (1983) demonstrated true endogenous adventitious buds developing *in situ* from dedifferentiated bark parenchyma, but also showed that exogenous axillary, collateral, or serial accessory buds may remain dormant and subsequently become engulfed within the bark during secondary growth. Such concealed buds can therefore appear endogenous despite their preventitious origin. Moreover, continuous vascular connection toward the pith is not an infallible diagnostic character. Fink found preventitious buds that secondarily lost their original vascular connection and adventitious buds that developed vascular strands toward traces of neighboring preventitious buds. He consequently concluded that developmental origin may ultimately require detailed ontogenetic analysis. If even internal vascular anatomy may incompletely preserve developmental history, the absence of separable classes in external shoot morphology is perhaps not unexpected.

The developmental behavior of epicormic buds provides a possible explanation for this morphological convergence. Fontaine et al. (1998) showed that proventitious epicormic buds of *Quercus petraea* do not contain a preformed shoot; rather, the shoot emerging from an activated bud is entirely neoformed by its terminal meristem. Although the same developmental process has not been demonstrated in the present *Prunus* stump, this observation provides a plausible model for the present result. Once shoot development begins, much of the visible phenotype may reflect subsequent shoot growth rather than the developmental history of the bud from which it arose. Different developmental pathways could therefore converge on a similar post-emergence growth program.

This interpretation distinguishes the present analysis from previous quantitative studies of epicormic and stump-sprout morphology. Fontaine et al. (2004) quantitatively classified epicormic formations of *Q. petraea* using external morphological characteristics and related these formations to traces and defects within the wood. Similarly, stump-sprout studies have assigned shoots to proventitious or adventitious categories using position, attachment, or other developmental criteria and subsequently measured their quantitative characteristics; Ríos-Saucedo et al. (2017) provides one example. The present study posed the inverse problem: developmental origin was left unlabeled, and the quantitative whole-shoot phenotype itself was tested for evidence of latent populations. The inability to relate the recovered morphological structure to developmental origin suggests that quantitative whole-shoot morphology alone may be poorly suited to retrospective inference of bud origin even though it remains informative for describing developmental state and sprouting vigor.

The occurrence of attached woody tissue initially appeared to provide an additional candidate marker of developmental origin. However, attached wood was strongly associated with shoot size: larger shoots were substantially more likely to retain woody tissue during collection. This dependence argues against treating attached wood as an independent developmental marker. It may instead reflect increasing mechanical integration of a growing shoot with the parent stump, differences in the fracture plane during removal, or both. These mechanisms were not directly tested. More generally, the result is consistent with the anatomical caution raised by Fink (1983): apparent structural relationships at the shoot–stem junction need not directly reveal developmental origin. In the present material, attached woody tissue therefore provides information about the physical shoot–stump junction but cannot by itself be used to classify a sprout as preventitious or adventitious.

A more unexpected observation was the basal longitudinal cavity (BLC). Among the sectioned sprouts from the initial census, the BLC was present in 23 of 31 individuals (74.2%), whereas it occurred in 16 of 22 sprouts (72.7%) in the second cohort. Thus, despite the substantial difference in overall shoot size between cohorts, BLC prevalence was essentially unchanged (Fisher’s exact test, *P* = 1.0). In contrast, BLC axial length differed between cohorts. Among BLC-positive sprouts, median BLC length was 3.9 mm in the initial cohort and 1.05 mm in the second cohort (*P* = 0.0086). Moreover, BLC length was positively associated with the common growth-size axis within each cohort independently (24 August: Spearman’s ρ = 0.452, *P* = 0.030; 31 August: ρ = 0.520, *P* = 0.039), as well as when the cohorts were pooled (ρ = 0.478, *P* = 0.0021). The combination of stable prevalence and growth-associated length suggests a possible distinction between **BLC establishment** and **BLC extension**: whether a cavity is present may be determined relatively early or independently of subsequent shoot growth, whereas its longitudinal extent may increase during shoot development. This interpretation remains descriptive because neither the tissue identity nor the mechanism producing the cavity was determined.

Internal spaces are known from developing woody buds and shoots, but the available precedents do not establish homology with the structure observed here. Sakai (1979) described internal organization associated with extraorgan freezing in conifer buds, while studies of *Prunus* flower buds have identified anatomical differences associated with contrasting freezing strategies (Kadir and Proebsting, 1994). Fink (1983) also described older endogenous adventitious buds of *Couroupita guianensis* in which free leaf primordia and outer scale-like organs occurred within a small “cavity.” These observations establish that internal spaces can occur as components of woody-bud development, but they concern different organs, taxa, and developmental contexts. The BLC observed in the present summer-growing stump sprouts should therefore not presently be interpreted as a pith cavity, a freezing adaptation, or a homolog of any previously described bud cavity. Rather, “basal longitudinal cavity” is used here as a purely descriptive term for the macroscopically visible longitudinal space extending from the basal end of the sprout.

Decay provides an alternative explanation because the parent stump was old, hollow, and visibly decayed. Roth and Hepting (1943) demonstrated that decay in a parent oak stump can progress into an attached sprout through their woody connection. Long-term observations of baldcypress stump sprouts similarly describe rot extending from old stumps into sprout bases (Keim et al., 2006). However, the second cohort places an important temporal constraint on a simple secondary-decay explanation. Twenty-two visible sprouts were collected only seven days after complete removal of the first cohort, yet BLC prevalence was almost identical to that of the initial population. It therefore seems difficult to explain the high frequency of BLCs in this cohort solely by progressive decay occurring within newly developed shoot tissue during those seven days. This observation does not exclude continuity with a pre-existing cavity or decayed region in the parent stump, nor does it establish a developmental origin for the BLC. Indeed, if a cavity or degraded region already existed within the underlying stump tissue, a newly emerging sprout might retain anatomical continuity with that structure. Histological continuity between the BLC and tissues of the stump therefore remains to be determined.

The second census also provides information about the regenerative organization of the stump itself. Classical work demonstrates that repeated sprouting need not arise through a single developmental pathway. In scrub oaks, Woods and Cassady (1961) found that initial stump sprouts were associated with dormant-bud traces, whereas removal of these sprouts was followed by callus formation and predominantly adventitious sprouting. Other species possess persistent basal or epicormic bud systems containing secondary or accessory buds capable of producing temporally separated shoots. Importantly, adventitious buds themselves may also persist in a dormant state after their formation. Consequently, the appearance of 22 sprouts within seven days of the first complete census cannot itself distinguish among activation of previously inactive preventitious buds, activation of pre-existing dormant adventitious buds, development of buds or shoots already initiated but not externally visible on 24 August, and de novo adventitious bud formation following removal.

The short interval nevertheless makes the second cohort particularly informative. Woods and Cassady (1961) reported that resprouting from callus began rapidly after sprout removal, but that at least two weeks were required for sprouting to develop from dormant buds in a freshly made scrub-oak stump. The appearance of numerous *Prunus* sprouts within seven days in the present study is therefore an interesting contrast, but this difference cannot be used to assign their origin because species, stump age, previous injury history, climate, and the developmental state of the underlying buds differ. More fundamentally, time since the most recent disturbance is not equivalent to bud age. A shoot appearing after experimental removal may originate from a bud whose developmental history substantially predates that removal. This distinction is particularly important when interpreting preventitious and adventitious origins, because “adventitious” describes developmental origin rather than the time at which a bud became visible or began elongating.

The reproducibility of BLC prevalence across the two censuses is noteworthy in this context. The first and second cohorts differed markedly in their overall quantitative morphology, yet approximately three quarters of the sectioned sprouts in each cohort contained a BLC. At the same time, BLC length was shorter in the second cohort and positively related to PC1 within each cohort. These observations are compatible with a feature that is established at or before an early stage of visible shoot development and subsequently extends as the shoot grows. Whether that early state reflects bud developmental origin, the organization of the shoot–stump junction, continuity with pre-existing parent tissue, or another process cannot be distinguished from the present data. Nevertheless, the reproducibility of the BLC contrasts with the failure of whole-shoot quantitative morphology to provide a basis for distinguishing developmental origins and identifies basal anatomy as a potentially more informative level at which to investigate developmental history.

Several limitations define what can be concluded from this study. First, developmental origin was not independently established by serial anatomical reconstruction or histology, so the presence or relative frequency of preventitious and adventitious sprouts remains unknown. The inability to relate the observed morphological structure to developmental origin therefore cannot be interpreted as evidence that only one developmental origin was present. Second, the study concerns a single, taxonomically unresolved ornamental cherry (*Prunus* sp.) stump. The complete-census design eliminates variation among source trees within the observed population and permits the entire visible sprout population of that stump to be examined, but it does not provide biological replication among trees. Third, the spatial positions of individual sprouts were not recorded because the highly irregular stump surface did not permit reproducible positional coordinates. Spatial clustering corresponding to individual underlying bud systems therefore cannot be reconstructed retrospectively. Finally, attached woody tissue and the BLC were recognized during examination of the collected material rather than being among the four quantitative traits selected before the first census. Analyses involving these basal features should therefore be regarded as exploratory rather than as tests of pre-specified anatomical hypotheses.

These limitations also identify the most informative next experiment. Serial longitudinal and transverse sections through the shoot–stump junction could determine whether individual sprouts possess traces continuous with primary xylem, arise from callus or other secondary tissues, or connect with pre-existing bud complexes. Histological examination could additionally identify the tissues bordering the BLC and determine whether the cavity is continuous with the parent stump, arises within the developing shoot, or represents tissue degradation. However, Fink’s observations caution that vascular continuity alone may not always provide an unambiguous developmental diagnosis: preventitious buds may lose or lack apparently continuous traces, whereas adventitious buds may acquire vascular continuity with older traces. Ideally, serial anatomy would therefore be combined with direct ontogenetic observations to provide independent developmental labels against which quantitative morphology and BLC anatomy could then be tested.

## Conclusion

Two complete censuses of stump sprouts from a single ornamental cherry showed that quantitative whole-shoot morphology was dominated by a reproducible common growth axis. Although unsupervised mixture modelling detected statistical components within the morphological distribution, these components could not be related to the hypothesized preventitious and adventitious developmental origins. Complete removal of the initial population was followed within seven days by a smaller second cohort that nevertheless reproduced the same coordinated morphological architecture. Thus, statistical structure in whole-shoot morphology did not provide a basis for distinguishing developmental origin. Most notably, the basal longitudinal cavity occurred at nearly identical frequencies in the initial and second cohorts despite their substantial difference in shoot size and the complete removal of the first cohort only seven days earlier. Whereas cavity occurrence was largely independent of overall shoot size, cavity length increased along the common growth-size axis among cavity-positive sprouts. This combination of reproducible occurrence and growth-associated extension suggests that the cavity represents a stable anatomical feature established relatively early in sprout development and subsequently extended during growth, although its developmental basis remains unknown. Thus, preventitious and adventitious origins could not be distinguished from gross quantitative morphology, but developmental history may still be reflected in basal anatomy. Direct anatomical and ontogenetic examination will be required to determine whether the basal longitudinal cavity is associated with developmental origin and to establish the origin of individual sprouts.

## Author contributions

NNO performed all stump-sprout collection and measurements. NO conceived and designed the study, performed all data analyses and visualization, interpreted the results, conducted the literature investigation, and wrote and revised the manuscript.

